# EISCA and EISTA: Full-Spectrum Pipelines for Single-Cell and Spatial Transcriptomics Analysis

**DOI:** 10.64898/2026.09.02.748851

**Authors:** Huihai Wu, Ashleigh Lister, Iain Macaulay, Katie Long, Cristobal Uauy, Yuxuan Lan, Gregory J. Wickham, David Swarbreck, Pavankumar Videm, Andrew Stubbs, Nicola Soranzo, Myrthe de Waard - van Baardwijk, Amirhossein Naghsh Nilchi, Irene Papatheodorou

**Author notes:** Correspondence to: Huihai Wu.

## Abstract

Single-cell and spatial transcriptomics are transforming our understanding of cellular heterogeneity and tissue organization, yet their analytical complexity remains a major bottleneck. Here, we present EISCA and EISTA, two standardized, end-to-end pipelines for single-cell RNA-seq and imaging-based spatial transcriptomics analysis. Built on the Nextflow nf-core framework, both pipelines implement modular, scalable, and reproducible workflows spanning primary, secondary, and tertiary analyses, from raw data processing to advanced downstream analyses. EISCA supports droplet- and plate-based scRNA-seq technologies, while EISTA is tailored for high-resolution spatial platforms including Vizgen MERFISH and 10x Xenium. Together, they integrate state-of-the-art methods for quality control, normalization, clustering, integration, cell-type annotation, differential expression, and cell-cell communication, with EISTA further enabling spatial statistical analyses. A central design principle is to balance standardization with flexibility: workflows can be executed end-to-end or modularly, enabling iterative, exploratory analyses with minimal overhead. Both pipelines deliver rapid preliminary results alongside an out-of-the-box report, facilitating immediate data assessment and accelerating downstream discovery. Case studies in plant immunity and human sepsis demonstrate that EISTA and EISCA reproducibly can be used to recover biologically meaningful insights. Collectively, these pipelines provide efficient, flexible, and scalable solutions for comprehensive single-cell and spatial transcriptomics analyses.

## 1 Introduction

Biological processes are executed by groups of cells that interact with each other within specific spatial microenvironments. The development of single-cell RNA sequencing (scRNA-seq) has transformed our ability to interrogate biological processes at cellular resolution, enabling the discovery of previously unrecognized cell types and generating mechanistic insights into development, homeostasis, and disease (Macosko et al., 2015; Zheng et al., 2017). For example, mapping of the tumor microenvironment has revealed prognostically relevant malignant programs and immune-cell dysfunction linked to immunotherapy response (Sade-Feldman et al., 2018). However, the dissociation required for scRNA-seq disrupts tissue architecture and erases spatial context. To overcome the loss of spatial information inherent to single-cell methods, spatial transcriptomics has emerged to map gene expression within intact tissue sections, linking molecular profiles to histology and microanatomy (Larsson et al., 2021; Ståhl et al., 2016). With these technologies, researchers can directly visualize and quantify spatial expression patterns and infer spatially resolved cell–cell interactions. For example, researchers combined single-cell and spatial transcriptomics to uncover how rice root cell types respond to soil compaction and nutrient heterogeneity, mapping cell-type–specific transcriptomic adaptations in situ (Zhu et al., 2025). Another study identified oncogenic characteristics shared by each tumor subclone, including T-cell and macrophage infiltration, immune activity, and antigen presentation, painting a detailed picture in both two- and three-dimensional space of how tumors evolve within their microenvironment (Mo et al., 2024). Despite the rapid progress in spatial transcriptomics, challenges remain, including trade-offs between resolution, sensitivity, and transcriptome breadth. Addressing these issues through methodological innovation and community standards will further enhance the power of single-cell and spatial transcriptomics to decode tissue organization, improve diagnosis and prognosis, and accelerate target discovery and precision therapeutics.

The rapid expansion of single-cell technologies has catalyzed a rich ecosystem of algorithms, toolkits, and pipelines spanning single-cell and spatial transcriptomics. The R-based Seurat package is a widely adopted end-to-end framework that supports rigorous quality control (QC), normalization, dimensionality reduction, clustering, differential expression, and data integration (Stuart et al., 2019). An alternative python package, Scanpy, builds on the AnnData data model to deliver similar functionalities (Wolf et al., 2018). Scanpy often exhibits great scalability on large datasets through sparse operations, easier integration with modern machine learning frameworks, and seamless coupling with GPU-enabled libraries. For instance, scvi-tools provides a PyTorch-based suite of deep generative models for probabilistic normalization, batch correction, multi-omic integration, and spatial deconvolution, enabling uncertainty-aware analyses at scale (Gayoso et al., 2022). For spatial transcriptomics specifically, Squidpy extends the Scanpy/AnnData ecosystem with spatial graphs, neighborhood enrichment analysis, spatial autocorrelation analysis, and image feature extraction alongside interactive visualization (Palla et al., 2022). Several pipelines have been developed to standardize single-cell workflows based on these packages. The nf-core/scrnaseq pipeline implements best practices for processing 10x scRNA-seq data from raw FASTQ files to count matrices with standardized QC and reporting within the Nextflow nf-core framework (Ewels et al., 2020). scDown (Sun et al., 2025) streamlines downstream single-cell analysis into a reproducible, report-driven workflow accessible to non-experts. Benchmarking studies using mixture control experiments have clarified how choices of normalization, feature selection, dimensionality reduction, and clustering affect sensitivity and specificity, guiding practitioners toward well-calibrated single-cell pipelines (Tian et al., 2019). In the spatial transcriptomics domain, nf-core/spatialvi standardizes Visium data processing and downstream modeling, integrating image-aware QC, mapping, and scvi-tools–based spatial latent variable models. The pipeline nf-core/spatialxe provides best-practice processing and quality control for imaging-based spatial transcriptomics data such as Vizgen MERFISH and 10x Xenium, focusing primarily on segmentation preprocessing. Another spatial analysis framework, Giotto, provides tools for preprocessing, visualization, spatial statistics, and cell–cell interaction analysis (Dries et al., 2021). In this work, we focus on in situ imaging–based technologies, such as Vizgen’s MERFISH, which use multiplexed fluorescent readouts of targeted probe panels to achieve subcellular resolution and single-molecule sensitivity in fresh-frozen or FFPE tissues (Chen et al., 2015).

We present two pipelines developed at the Earlham Institute: EISCA (Earlham Institute Single-Cell Analysis) for scRNA-seq and EISTA (Earlham Institute Spatial Transcriptomic Analysis) for spatial transcriptomics. Both pipelines encompass primary, secondary, and tertiary analysis phases and are implemented using the Nextflow framework, following nf-core best practices, and the Scanpy/AnnData ecosystem to provide generalizable, flexible, and scalable workflows. EISCA (github.com/EarlhamInst/eisca) supports both droplet-based platforms such as 10x Chromium and plate-based technologies such as Smart-seq2, while EISTA (github.com/EarlhamInst/eista) currently supports Vizgen MERFISH and 10x Xenium technologies. Single-cell analysis is inherently diverse, dynamic, and exploratory; therefore, our objective is to develop pipelines that not only enable robust end-to-end analysis but also retain exploratory capabilities across different analyses. Ultimately, we aim to develop pipelines that are standardized, scalable, efficient, and flexible. The pipelines offer four key advantages: 1) ease of use—no manual tool installation is required, and execution is supported on local machines, HPC clusters, and cloud platforms; 2) flexibility—pipelines can be run end-to-end or by module, with parameters adjustable to specific analyses; 3) standardization—workflows rely on widely used Python packages to process large datasets consistently and efficiently; and 4) extensibility—the pipelines provide a robust foundation for routine analyses while enabling integration of task-specific modules. Both pipelines can rapidly generate preliminary results together with out-of-the-box reports, facilitating initial data assessment and a smooth transition to advanced downstream analyses. To our knowledge, EISTA is one of the few end-to-end pipelines available for spatial transcriptomics analysis.

## 2 Materials and methods

### 2.1 Overview of pipelines

EISCA is a pipeline designed to process droplet-based (10x scRNA-seq) and plate-based (Smart-seq2) single-cell transcriptomic data. The metro map in Figure 1 illustrates the workflow and modules of EISCA. The primary analysis starts with raw sequencing data and proceeds through sequence quality control, alignment, quantification, and count matrix generation, ultimately producing raw counts. The secondary analysis focuses on cell-level quality control, filtering, normalization, and clustering, resulting in processed and annotated counts suitable for downstream analyses. The tertiary analysis covers common downstream analyses, including cell-type annotation using CellTypist or scvi-tools, differential expression analysis (DEA) with Scanpy or scvi-tools, and cell–cell interaction analysis using CellChat.

**Figure 1.**
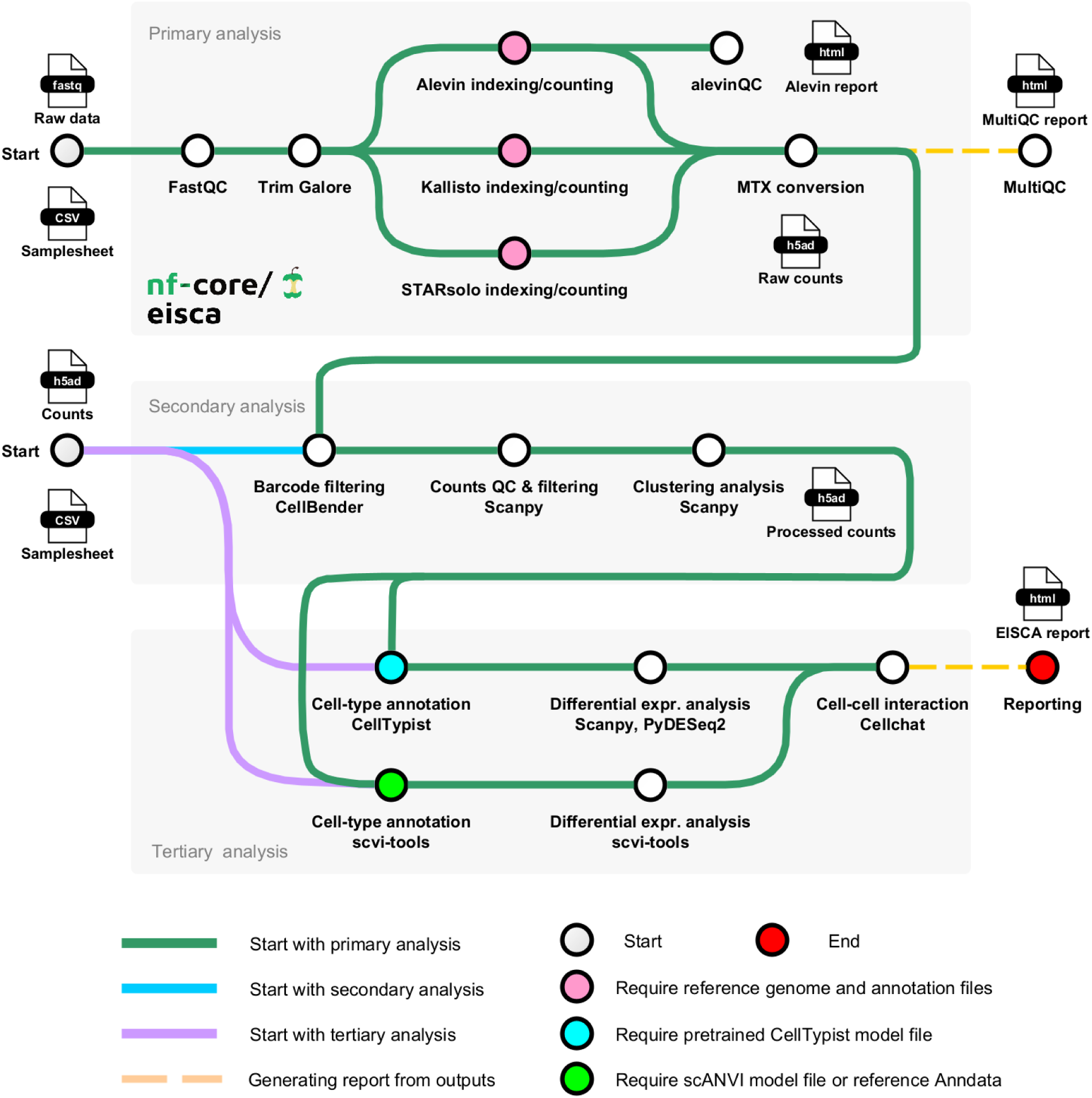
Metro map illustrating the EISCA workflow for single-cell transcriptomic data.

EISTA is a bioinformatics pipeline designed for the analysis of single-cell spatial transcriptomics data. Figure 2 shows a metro map illustrating the EISTA workflow and its modules. The EISTA pipeline is primarily designed for Vizgen MERFISH data and also supports 10x Xenium data. In the primary analysis phase, Vizgen post-processing and count generation are performed for Vizgen imaging data, or Xenium data are read into count matrices. The secondary analysis includes cell-level quality control, filtering, clustering, and spatial statistical analysis. The tertiary analysis covers common downstream tasks similar to those in EISCA, extended with spatial mapping.

**Figure 2.**
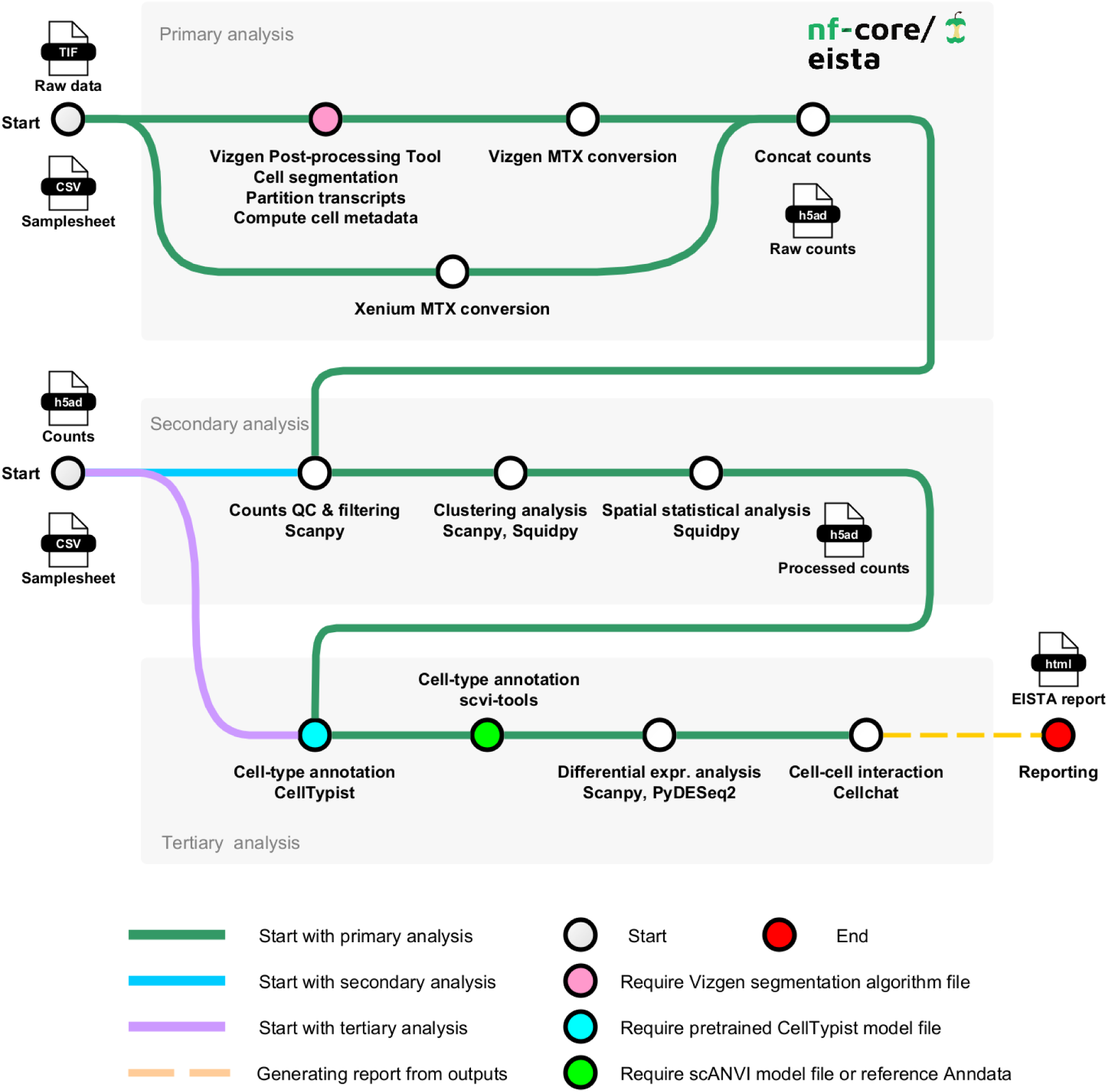
Metro map illustrating the EISTA workflow for spatial transcriptomics data.

Both pipelines can be executed starting either from raw data or from a pre-generated count matrix. The workflow can be run end-to-end, phase by phase, or for specific analyses. Figure 3 shows a typical exploratory analysis workflow using EISTA, from cell segmentation to spatial statistical analysis.

**Figure 3.**
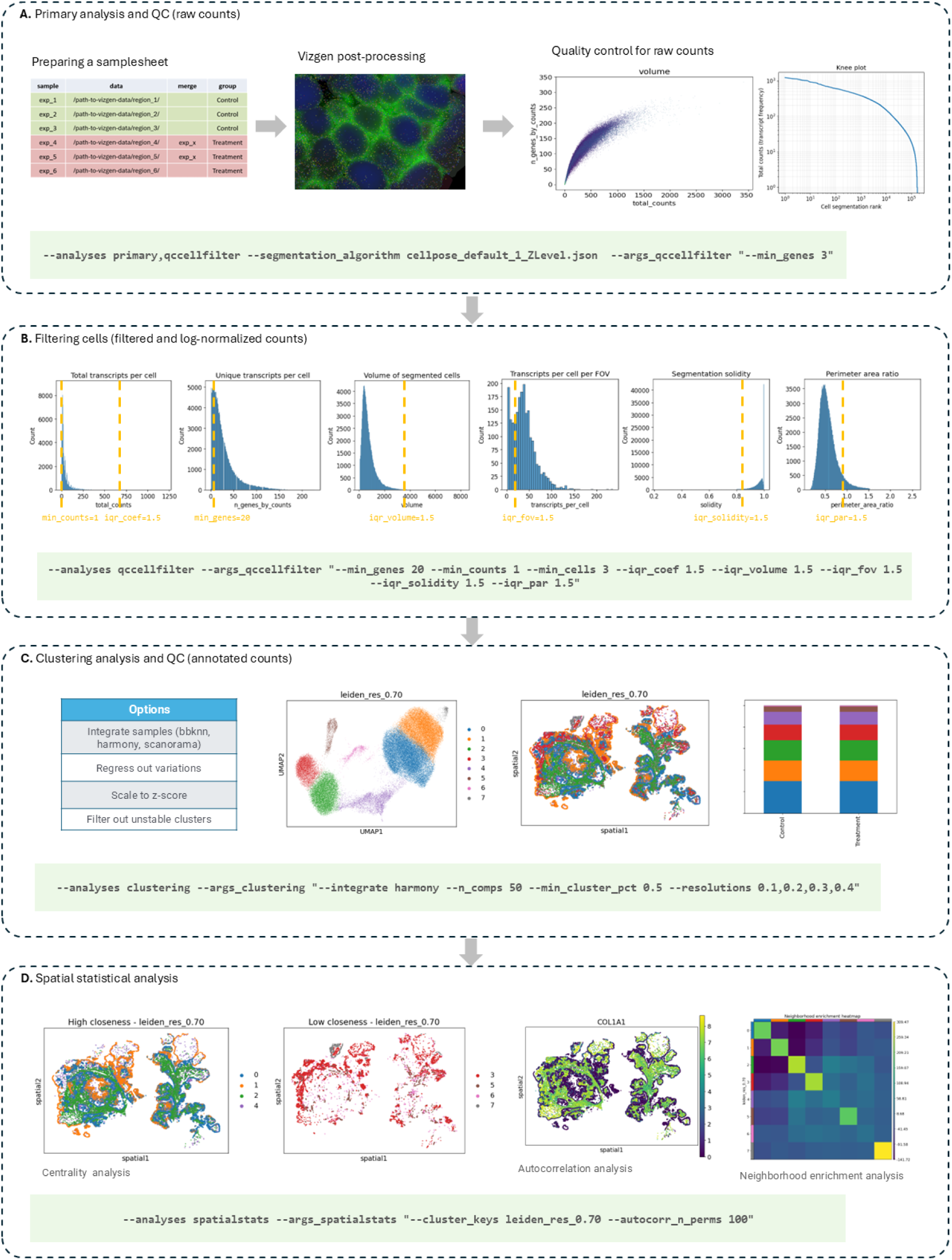
Schematic overview of the exploratory analysis workflow using EISTA. **A.** Preparing a sample sheet to define sample names, raw data locations, groups, and merging. Then applying VPT to obtain segmented cells, counts, and spatial metadata. Here the visualization shows segmented cells, and the scatter plot and knee plot illustrates count quality. **B.** Filtering cells to improve the accuracy of downstream analysis, here shows histogram of quality metrics and thresholds marked yellow. This step allows us to assess the raw count quality and adjust filtering parameters accordingly. **C.** Clustering with data integration to identify reliable subpopulations, here shows a clustering UMAP, its spatial scatter plot and cluster proportions. This step helps identify cell types and provides an additional layer of quality control. **D.** Calculating spatial statistics to reveal the relationship between expression patterns and biological morphology. The spatial scatter plots at left illustrate that clusters with high and low closeness centrality; the spatial scatter plot of gene COL1A, showing strong spatial autocorrelation and indicating that it is a spatially variable gene; the heatmap shows the results of the neighborhood enrichment analysis indicating that clusters are primarily self-enriched with mild overlaps, where the diagonal represents self-enrichment.

### 2.2 Primary analysis of EISCA

The primary analysis focuses on converting raw sequencing data into a gene–cell count matrix. This process includes raw read quality control, mapping/alignment, quantification, and count matrix generation. The inputs to the primary analysis are the raw sequencing data and a sample sheet that defines sample names, corresponding data file locations, samples to be merged, and group assignments. This metadata is essential for downstream analyses.

First, the quality of raw scRNA-seq data can be assessed using FastQC (Andrews, S. (2010), which provides an overview of potential issues that should be addressed prior to downstream processing. If a portion of the sequencing data shows poor quality (Phred quality score < 20), trimming can be applied to the FASTQ files using TrimGalore (Martin, 2011), which can also remove sequencing adapters. Although adapter and base trimming is usually unnecessary for 10x droplet-based data, it is recommended for plate-based protocols such as Smart-seq2, which is similar to bulk RNA-seq. Adapter contamination can be identified by inspecting the overrepresented sequences section of the FastQC report.

The next step is mapping/alignment and quantification, for which three toolsets are supported: Kallisto & Bustools (Sullivan et al., 2025), Salmon Alevin (Patro et al., 2017), and STARsolo (Kaminow et al., 2021). The goal of this step is to estimate the number of RNA molecules per gene per cell, which represents the gene expression profile of each cell. We recommend Kallisto & Bustools or Salmon Alevin for large-scale datasets, as these pseudo-alignment-based methods focus on gene-level quantification rather than determining exact read alignment positions. Consequently, they are substantially faster and require less memory than STARsolo. However, STARsolo is a splice-aware aligner and should be preferred when downstream analyses require splice-aware information, such as alternative splicing or RNA velocity analysis. The output of this step is a raw count matrix, which is then converted and combined into an AnnData object stored in H5AD format. This object stores the expression matrix together with cell-level and gene-level metadata, additional data layers, and other analysis-related information, and serves as the input for quality control and filtering in the secondary analysis.

### 2.3 Primary analysis of EISTA

The primary analysis focuses on converting imaging-based spatial transcriptomics data into a count matrix. The Vizgen Postprocessing Tool (VPT) (Vizgen, 2026) is applied for cell segmentation, transcript partitioning, and computation of cell metadata. This step generates segmented cells, gene counts, and spatial metadata, which are subsequently converted into a raw count matrix (Figure 3A).

The Vizgen MERSCOPE instrument generates many output files. For EISTA, the minimal required inputs are organized in a directory structure where each sample has its own folder. Within each sample folder, at least four files are required: file *detected_transcripts.csv* containing the decoded gene names together with their 3D spatial coordinates; under the subfolder *images*, file *mosaic_DAPI_z3.tif* storing the stitched DAPI nuclear staining image at Z-plane 3, while *mosaic_PolyT_z3.tif* storing the stitched PolyT cytoplasmic staining image at the same Z-plane; file *micron_to_mosaic_pixel_transform.csv* providing the coordinate transformation used to convert physical micron coordinates into mosaic image pixel coordinates. To provide sample-level metadata for the pipeline, a samplesheet file must also be prepared. This file defines the sample names, the samples to be merged, the corresponding full paths to the Vizgen data for each sample, and the assigned groups. The column *group* is required for most downstream analyses, as it defines the experimental conditions used for comparisons.

For Vizgen MERFISH data, we applied VPT for image preprocessing and quantification. VPT is a toolset for imaging data processing, quantification, and visualization. In this module, five main subprocesses were performed: 1) Cell segmentation – a segmentation algorithm is applied to individual tiles of the mosaic images, and per-tile segmentation results are merged into a single internally consistent parquet file containing all cell boundary definitions. 2) Transcript partitioning – detected transcripts are assigned to segmented cells using the computed cell boundaries. 3) Cell metadata calculation – geometric properties of each segmented cell are computed from the segmentation boundaries. 4) Signal summarization – image intensity values are aggregated within each segmented cell to quantify transcript signals. 5) VZG file updating – the existing VZG file can be updated with refined segmentation boundaries and the corresponding expression matrix.

The segmentation process in the primary analysis can also be customized by specifying a Cellpose algorithm JSON configuration file. By default, the primary analysis uses the configuration file *cellpose_default_1_ZLevel.json* stored in the EISTA pipeline. This configuration file allows the specification of either a single Z-plane or multiple Z-planes using the *z_layers* option. Z-plane 3 is usually the central focal plane and is often sufficient for cell segmentation. The output of the primary analysis is a raw gene–cell count matrix in a AnnData object, saved as an H5AD file for downstream analyses.

### 2.4 Secondary analysis

Secondary analysis focuses on preprocessing raw count data to maximize the signal-to-noise ratio of gene expression profiles and ensure comparability across samples. The main tasks include quality control, cell filtering, clustering analysis, and sample integration. For EISTA, spatial statistical analysis is also included in the secondary analysis, as it reveals relationships between expression patterns and tissue morphology and provides spatial quality control of the data. Analyses are primarily implemented using the Python package Scanpy (Wolf et al., 2018) and the additional package Squidpy (Palla et al., 2022) for EISTA.

The quality of raw count data can be assessed using several diagnostic plots (Figure 3A). The scatter plot illustrates count quality by showing the relationship between total counts and the number of detected genes. For a high-quality dataset, a saturating curve is typically observed, characterized by a steep rise at lower counts that levels off at higher counts. The knee plot shows the relationship between cell barcode/segmentation rank and the total number of transcripts per barcode/segmentation. For a high-quality dataset, the curve remains relatively flat before showing a sharp drop toward the end of the range. EISCA can generate violin plots summarizing the distributions of cells by the number of detected genes, total counts, and the percentage of mitochondrial gene counts. EISTA can generate histogram plots summarizing the distributions of cells by the number of detected genes, total counts, and other segmentation metrics. These distribution plots provide useful guidance for quality assessment and cell filtering.

Cell filtering is a critical step in single-cell RNA-seq analysis and involves removing low-quality or unwanted cells to improve the accuracy of downstream analyses and the reliability of biological conclusions. Both pipelines provide multiple cell-filtering strategies, including hard thresholding, quantile-based filtering, and outlier filtering based on interquartile range (IQR) rules. For example, users may define lower thresholds to exclude damaged or low-complexity cells and upper thresholds to remove potential multiplets, while outliers can be identified using IQR rules on the distributions of metrics. For example, based on the distributions of counts and segmentation metrics of EISTA, appropriate filtering thresholds can be applied (Figure 3B), such as a minimum number of counts, an upper bound for total counts, and a minimum number of detected genes. Additional criteria may include an upper bound for cell volume, a lower bound for transcripts per cell per FOV, a lower bound for solidity, and an upper bound for the perimeter-to-area ratio. In EISCA, users can apply CellBender to remove empty droplets from droplet-based single-cell RNA-seq data based on a deep learning framework of variational autoencoders (VAEs) (Fleming et al., 2023). Users may also optionally perform doublet detection using Scrublet (Wolock et al., 2019), which outputs both predicted doublet labels and doublet scores. The output of the cell-filtering step is a filtered and log-normalized count matrix stored in H5AD format.

Clustering analysis groups cells into subpopulations based on similarities in their gene expression profiles (Figure 3C). Main steps in clustering analysis include: 1) identifying highly variable genes; 2) performing dimensionality reduction using Principal Component Analysis (PCA) and constructing a nearest neighbor graph; 3) applying the Leiden graph-clustering method. Visualization of clustering results aids cell-type identification and provides an additional layer of quality control, allowing users to further filter tiny or unstable clusters. Users may optionally apply additional preprocessing steps to improve clustering performance, including sample integration, removal of predicted doublets, regression of unwanted sources of variation (e.g., total counts per cell), and z-score transformation. Multiple clustering resolutions can be specified simultaneously, enabling systematic exploration of cluster granularity and facilitating the identification of biologically meaningful cell types. With spatial mappings of clusters, EISTA can further illustrate how transcriptionally defined clusters are organized across tissue morphology.

For multi-sample datasets, data integration is essential to correct batch effects arising from inter-sample or inter-experimental variability, such as differences in sample preparation, sequencing technology, or other technical factors. Both pipelines support three integration methods: BBKNN (Polański et al., 2020), a fast approach that corrects batch effects while preserving local neighborhood structure; Harmony (Korsunsky et al., 2019), a widely used global correction method that iteratively adjusts low-dimensional embeddings; and Scanorama (Hie et al., 2024), which emphasizes global dataset merging while retaining subtle biological variation across conditions. EISCA also supports integration with scvi-tools (Gayoso et al., 2021), which uses a deep generative model based on a conditional variational autoencoder to learn a latent representation that accounts for batch effects while preserving biological signals, making it particularly suitable for highly heterogeneous datasets. The output of the clustering and integration steps is an integrated low-dimensional representation of the cells, together with associated clustering metadata, saved in H5AD format. However, for imaging-based spatial transcriptomics datasets, integration should be applied cautiously. For example, in datasets with strong biological gradients, such as infection time courses, integration may be unnecessary and can lead to over-correction, potentially removing part of the biological effects (Figure S2).

Spatial statistical analysis of EISTA aims to characterize relationships between gene expression patterns and tissue architecture. This includes centrality analysis, neighborhood enrichment analysis, and spatial autocorrelation analysis (Figure 3D). For centrality analysis, three centrality scores are calculated to reflect how cell types are spatially organized and interact within the tissue architecture. For example, clusters with high closeness centrality act as global bridges between distant regions and tend to display a more dispersed distribution throughout the tissue, whereas clusters with low closeness centrality exhibit more isolated spatial patterns. Clusters with high degree centrality act as local hubs connecting to many neighboring clusters, whereas clusters with low degree centrality are often lower-abundance populations. A higher clustering coefficient indicates a stronger tendency for cells of the same cluster to group together locally, whereas a lower clustering coefficient suggests a more even distribution across the tissue. Heatmaps from neighborhood enrichment analysis indicate whether certain cell types are spatially enriched in neighboring regions of the tissue. A strong positive score means two clusters frequently appear as neighbors, whereas a strong negative score indicates spatial avoidance. For spatial autocorrelation analysis, Moran’s I statistic is used to evaluate whether genes exhibit spatial clustering or a random distribution across the tissue. Genes with significant Moran’s I values (I > 0.4 and q-value < 0.05) can be considered spatially variable genes. Spatial statistical analysis can therefore be used to assess whether the identified clusters exhibit meaningful spatial organization and capture aspects of the tissue’s spatial heterogeneity.

### 2.5 Tertiary analysis

Tertiary analysis focuses on downstream analyses performed on preprocessed count data, including cell-type annotation, differential expression analysis (DEA), and cell-cell communication analysis.

Cell-type annotation is a central task in single-cell analysis that defines the cellular composition of a dataset. Users can perform automated annotation using CellTypist (Xu et al., 2023), which applies logistic regression classifiers optimized by stochastic gradient descent. The inputs to CellTypist are a preprocessed AnnData object and a pretrained CellTypist model; 52 built-in models are available from the CellTypist repository. Alternatively, users may train a custom model using a manually annotated reference AnnData object. CellTypist supports both single-label and multi-label classification, and also has a majority-voting strategy based on local over-clustering to refine predictions. The output is an AnnData object containing predicted cell-type labels and associated confidence scores when majority voting is applied.

Alternatively, users may perform cell-type annotation with scvi-tools (Gayoso et al., 2021) by training a scANVI model on reference data or by applying a pretrained model. As scvi-tools is based on a variational autoencoder framework, it enables transfer learning by projecting query data into a pre-trained scANVI latent space while correcting for technical batch effects, allowing the model to assign query labels based on the reference’s established cell-type probability distributions. In contrast, CellTypist applies a fixed linear classifier to normalized expression profiles and therefore requires prior batch harmonization and close matching between reference and query data. Consequently, scvi-tools is better suited for cross-study or atlas-scale annotation tasks across heterogeneous datasets, whereas CellTypist provides a lightweight and interpretable solution for rapid annotation of well-matched datasets.

For single-cell data, differential expression analysis (DEA) can be performed at the cell level to identify genes whose expression differs significantly between cell subpopulations and to estimate log fold changes as effect sizes. Users may apply Scanpy’s *rank_genes_groups* function to perform DEA, with configurable options to identify cluster marker genes, compare conditions across all cells or within specific cell types. The top marker genes can typically be selected using thresholds such as log fold change > 5, score > 100 (t-test), and adjusted p-value < 0.05. In addition, robust marker genes should also stand out in visualizations, such as rank plots showing relatively high scores and dot plots indicating strong expression levels and a high percentage of expressing cells. DEA can also be performed at the sample level using PyDESeq2 (Muzellec et al., 2023), which identifies differentially expressed (DE) genes between groups by aggregating cell-level expression data into pseudo-bulk samples. For EISCA, users may alternatively perform DEA using scvi-tools, which implements a deep generative Bayesian framework that explicitly models single-cell count distributions and batch covariates. This approach enables more robust differential expression testing in the presence of dropout noise and technical variability and provides more reliable effect size estimates than rank-based tests, while Scanpy’s method remains advantageous for rapid exploratory analysis.

Cell-cell communication analysis aims to infer and quantify signaling interactions between cell groups from single-cell transcriptomic data, thereby revealing how cells coordinate biological processes through ligand-receptor signaling networks. We apply CellChat (Jin et al., 2021) to identify overexpressed ligands and receptors in each cell group based on a curated database of ligand-receptor interactions. The resulting models enable users to identify, analyze, and visualize intercellular communication networks. For example, aggregated communication network plots summarize the number of interactions or total interaction strength between cell groups across the whole network or within specific signaling pathways. Bubble plots display significant ligand-receptor pairs mediating communication between cell groups, illustrating how intercellular signaling is orchestrated through multiple molecular interactions.

### 2.6 Galaxy Implementation of EISTA

An EISTA-inspired Galaxy workflow has been developed by the European Galaxy team (https://galaxyproject.org/eu/people/), as part of the “Spatial2Galaxy” project (https://elixir-europe.org/how-we-work/scientific-programme/science/cmr/spatial2) within the “Cellular and Molecular Research” ELIXIR Commissioned Service. The Galaxy implementation of EISTA generally adheres to the original workflow, though it incorporates several specific modifications. By utilizing SpatialData objects as input, the workflow is compatible with both sequence-based and image-based spatial transcriptomics data. To facilitate this flexibility, certain preprocessing tasks, such as the VPT-based processing of Vizgen data, were removed from the primary workflow; however, the workflow remains compatible with SpatialData objects generated via any external preprocessing tool. Furthermore, the downstream analysis has been enhanced by replacing the CellChat tool with LIANA+ (https://doi.org/10.1038/s41556-024-01469-w) that aggregates ligand-receptor scores from multiple methods, including CellChat.

## 3 Case study with EISTA

To showcase the exploratory analysis workflow with EISTA, a Vizgen MERFISH dataset of *Arabidopsis thaliana* leaves was used. The dataset originates from the study (Nobori et al., 2025) investigating cellular immune responses in *A. thaliana* leaves following infection with *Pseudomonas syringae* pv. *tomato* DC3000 expressing an immune-triggering effector AvrRpt2 (hereafter *Pto* AvrRpt2). MERFISH experiments targeting 500 genes were performed on five samples: a mock control and four post-infection time points at 4, 6, 9, and 24 hours (denoted as 4 h, 6 h, 9 h, and 24 h). We conducted the analysis using EISTA following an exploratory workflow. First, we performed primary analysis, including QC and cell filtering. Next, we carried out clustering to obtain a cluster-annotated and filtered dataset for downstream analyses, including spatial statistical analysis, differential expression analysis, and label transfer for cell-type annotation.

We used Z-plane 3 imaging data for cell segmentation with the default Cellpose algorithm configuration, and raw counts were generated during primary analysis. The raw count quality appeared satisfactory based on count distributions and QC plots. To remove low-quality cells, we applied multiple hard cutoffs, including minimum gene counts, minimum total counts, and gene filtering based on the number of expressing cells. In addition, we applied distribution-aware filtering using IQR-based rules, including upper bounds for total counts and cell volume, and lower bounds for transcripts per cell per FOV and solidity, as well as an upper bound for the perimeter-to-area ratio (Figure 3B). After filtering, a substantial number of low-quality cells were removed. QC metrics showed improved distributions, and median gene counts per cell were reasonable across samples (Figure S1). The UMAP in Figure 4A shows clustering of normalized counts after filtering, colored by sample. A continuous gradient structure is observed rather than distinct sample-specific clusters, with the majority of samples overlapping but exhibiting a directional shift.

**Figure 4.**
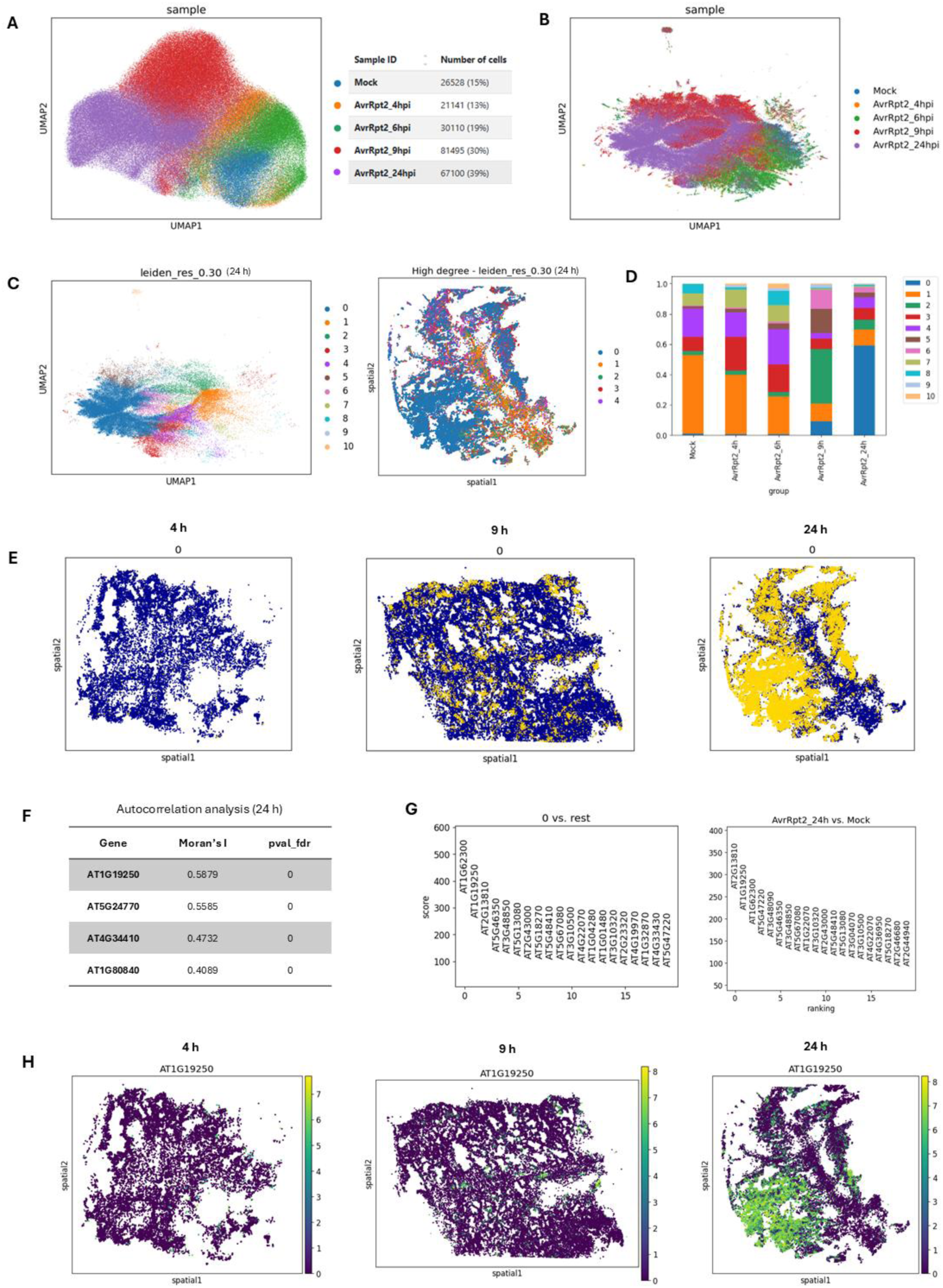
EISTA analysis on MERFISH dataset of Arabidopsis thaliana leaves. **A.** UMAP projection of filtered, normalized counts colored by sample, with a table indicating the number of cells remaining after filtering. **B.** UMAP projection of combined counts without integration after filtering out unstable clusters, colored by sample. **C.** Left: Leiden clustering UMAP at a resolution of 0.3 for the 24 h post-infection sample. Right: spatial mapping of high-degree clusters (0, 1, 2, 3, and 4), identified through spatial statistical analysis, revealing a structured spatial pattern associated with the infection response. **D.** This stacked bar plot compares cluster proportions across the five samples, with cluster 0 identified as the immune-active cluster. **E.** These are the spatial maps of immune-active cluster 0 for the three post-infection samples at 4 h, 9 h, and 24 h; the immune-active cells become more concentrated over time and are most abundant at 24 h. **F.** This table shows the top four spatially variable genes (Moran’s I > 0.4 and pval_fdr < 0.05) identified by autocorrelation analysis within the spatial statistical analysis module. **G.** Left: ranking plot of marker genes identified for cluster 0 from the differential expression analysis module. Right: ranking plot of differentially expressed genes in the 24 h sample compared with the mock control at the cellular level. And the gene AT1G19250 (FMO1) is known as a critical immune defense gene. **H.** These spatial maps highlight the expression of gene AT1G19250 in the three post-infection samples at 4 h, 9 h, and 24 h; clearly, the gene is most highly expressed in the 24 h sample.

Clustering analysis was performed on the filtered, log-normalized count matrix. Initial clustering revealed numerous small satellite clusters surrounding major clusters. These clusters appeared across samples and persisted across resolutions, suggesting technical artifacts such as segmentation fragments that were not sufficiently removed during the initial QC. We removed them by filtering clusters below a minimum proportion threshold of 0.5%. After removal, the UMAP structure became clearer, with eleven major clusters remaining (Figure 4C), likely representing true biological population. The five clusters (0–4) mapped onto the tissue (Figure 4C) showed high degree centrality in the spatial statistics analysis, with clusters 0, 1, 2, and 3 displaying concentrated spatial structures. Notably, cluster 0 is dominant in the 24 h sample (Figure 4D), and the inspection of cluster 0 in each time point (Figure 4E), implying it represents immune-active cells.

We further investigated cluster 0 using spatial statistical and differential expression analyses. The autocorrelation analysis results (Figure 4F) use Moran’s I to assess spatial structure. Four genes show significant spatial autocorrelation (Moran’s I > 0.4 and FDR-adjusted p < 0.05), indicating spatially variable genes that may serve as markers of tissue organization or infection response. To further examine potential maker genes, differential expression analysis was performed for both cluster marker identification and condition comparison. DEA results in Figure 4G reveal top three marker genes (log-foldchange > 5, score > 200, adjusted p-value < 0.05), AT1G19250, AT2G13810, and AT1G62300 in cluster 0, which are also upregulated in the 24 h sample relative to mock. All three genes are associated with immune responses in *A. thaliana*. AT1G19250 and AT2G13810 encode FMO1 and ALD1 respectively, key enzymes required for systemic acquired resistance (SAR) (Hartmann et al., 2018). AT1G62300 encodes the immunity-related transcription factor WRKY6 (Yang et al., 2024). These results demonstrate EISTA’s capability to identify immune-related genes exhibiting spatially structured expression in the combined MERFISH data.

Finally, we performed cell-type annotation by label transfer using scvi-tools, which enables transfer learning from an annotated snRNA-seq dataset to predict cell types in the Vizgen MERFISH data by training a scANVI model. The reference dataset, generated in the same study, contains three major annotated cell types: epidermis, mesophyll, and vasculature identified across the infection time course. Epidermis and mesophyll cells were further classified as immune-active or non-immune-active based on clustering analysis and defense marker gene expression. By combining cell-type identity with immune state, we defined five labels for downstream analysis: Epidermis_immune-active, Epidermis_non-immune-active, Mesophyll_immune-active, Mesophyll_non-immune-active, and Vasculature. The left UMAP in Figure 5A shows the predicted labels projected onto the scANVI latent space for the 24 h sample. The right UMAP shows the corresponding *scanvi_prob* values, representing the posterior probability that each cell belongs to its assigned label and therefore the confidence of the label transfer. Overall, the predictions showed high confidence, and a minimum threshold of 0.8 was applied to retain highly reliable predictions for downstream visualization. Figure 5B shows a stacked bar plot of the proportions of these high-confidence predicted labels across the five samples. Immune-active epidermis and mesophyll cells increased in proportion over the infection time course, with Mesophyll_immune-active cells becoming the most abundant population at 24 h. The spatial maps of predicted labels in Figure 5C show that immune-active epidermis and mesophyll cells progressively increase and become more spatially concentrated over the post-infection time course, with the strongest immune-active signal observed at 24 h and most dominated by immune-active mesophyll cells. In contrast, vasculature-associated cells show a more moderate change over time. These results are consistent with the observations reported in the original study and recover the expected temporal expansion of immune-responsive cell populations. These results suggest that scANVI-based label transfer is useful for assigning tissue-level labels across experiments. However, the approach is limited by multiple factors. For instance, the MERFISH dataset contains a limited gene panel; the reference annotation may contain uncertainty; sample cells may include background stress-like states; and label transfer assigns cells to the nearest available reference labels which have similar transcriptional profile. Therefore, the label transfer results should be interpreted as broad tissue-context annotations that reveal spatial immune activation, rather than as a fully resolved quantitative separation of all cell states.

**Figure 5.**
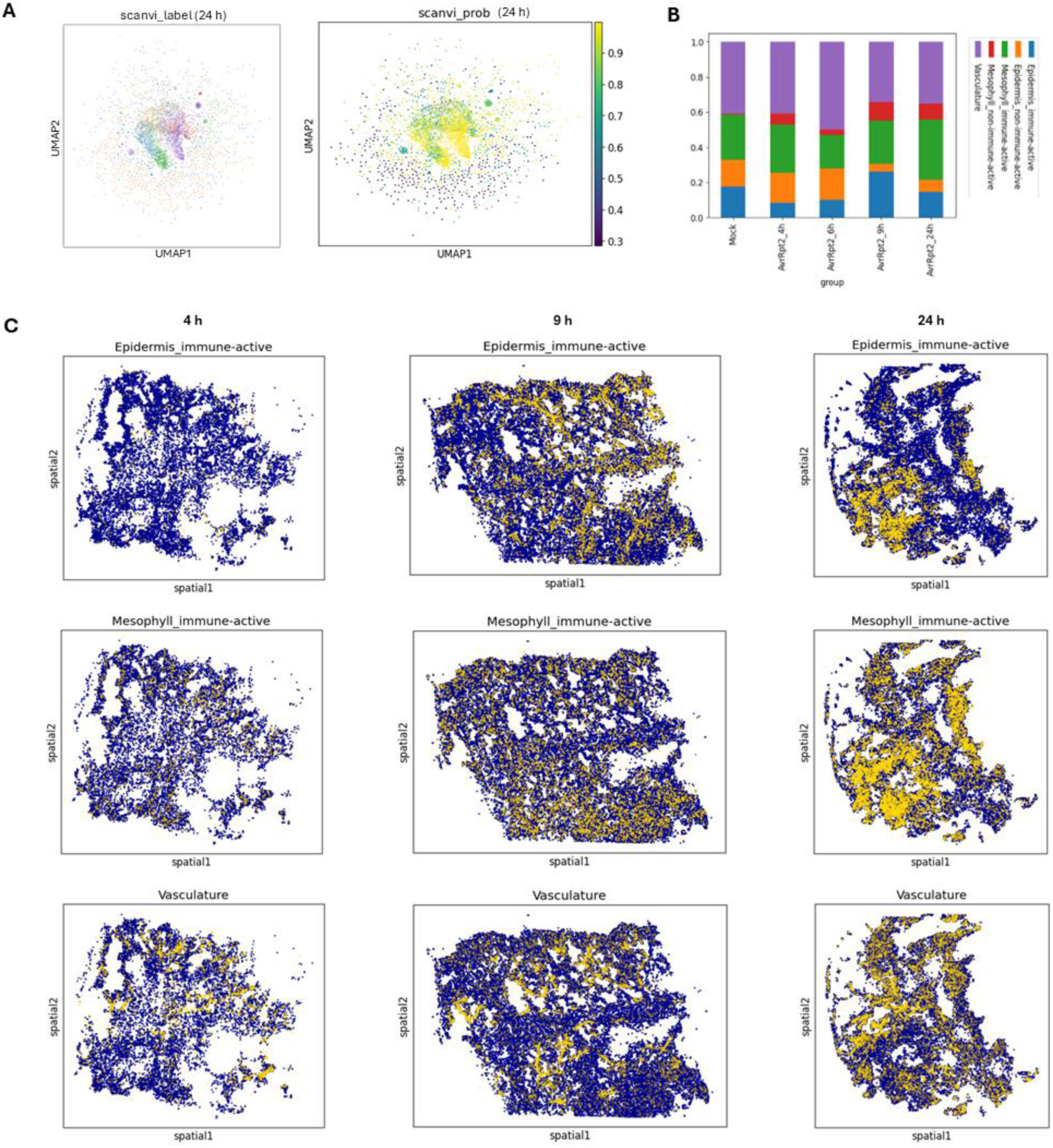
EISTA results on Vizgen MERFISH dataset of Arabidopsis thaliana leaves for cell-type annotation via label transfer from an immune-active snRNA-seq dataset using scvi-tools. **A.** Left: UMAP projection of predicted labels in the scANVI latent space, including immune-active and non-immune-active epidermis, mesophyll cells, and vasculature cells for the post-infection 24 h sample. Right: UMAP projection of predicted posterior probabilities (scanvi_prob) for assigned labels, indicating label transfer confidence. **B.** Stacked bar plot of predicted label proportions across five samples for cells with scANVI posterior probabilities greater than 0.8. **C.** These spatial maps show immune-active epidermis and mesophyll cells, and vasculature cells in the three post-infection samples at 4 h, 9 h, and 24 h.

## 4 Case study with EISCA

We applied EISCA to a set of scRNA-seq data from a human immune disorder sepsis study (Qiu et al., 2021), which conducted a comprehensive transcriptomic assessment of peripheral blood mononuclear cells (PBMCs) from a cohort of 12 samples comprising healthy controls (HC), sepsis survivors (SV), and non-survivors categorized by early-stage (NSES) and late-stage (NSLS) clinical outcomes (GSE167363). The analysis was performed from phase to phase. In the primary phase, raw sequencing data quality was examined, and counts were derived using Salmon Alevin. In the secondary phase, raw count quality was assessed, and a filtering process was performed to remove low-complexity cells based on gene/UMI thresholds and mitochondrial content (Figure 6A). Dimensionality reduction and Leiden graph clustering (evaluated across multiple resolutions) then produced stable clusters that were robust to parameter variation. Harmony-based integration mitigated inter-sample effects while preserving biologically meaningful structure, as evidenced by coherent UMAP separation of major immune lineages and consistent cluster composition across replicates (Figure 6B).

**Figure 6.**
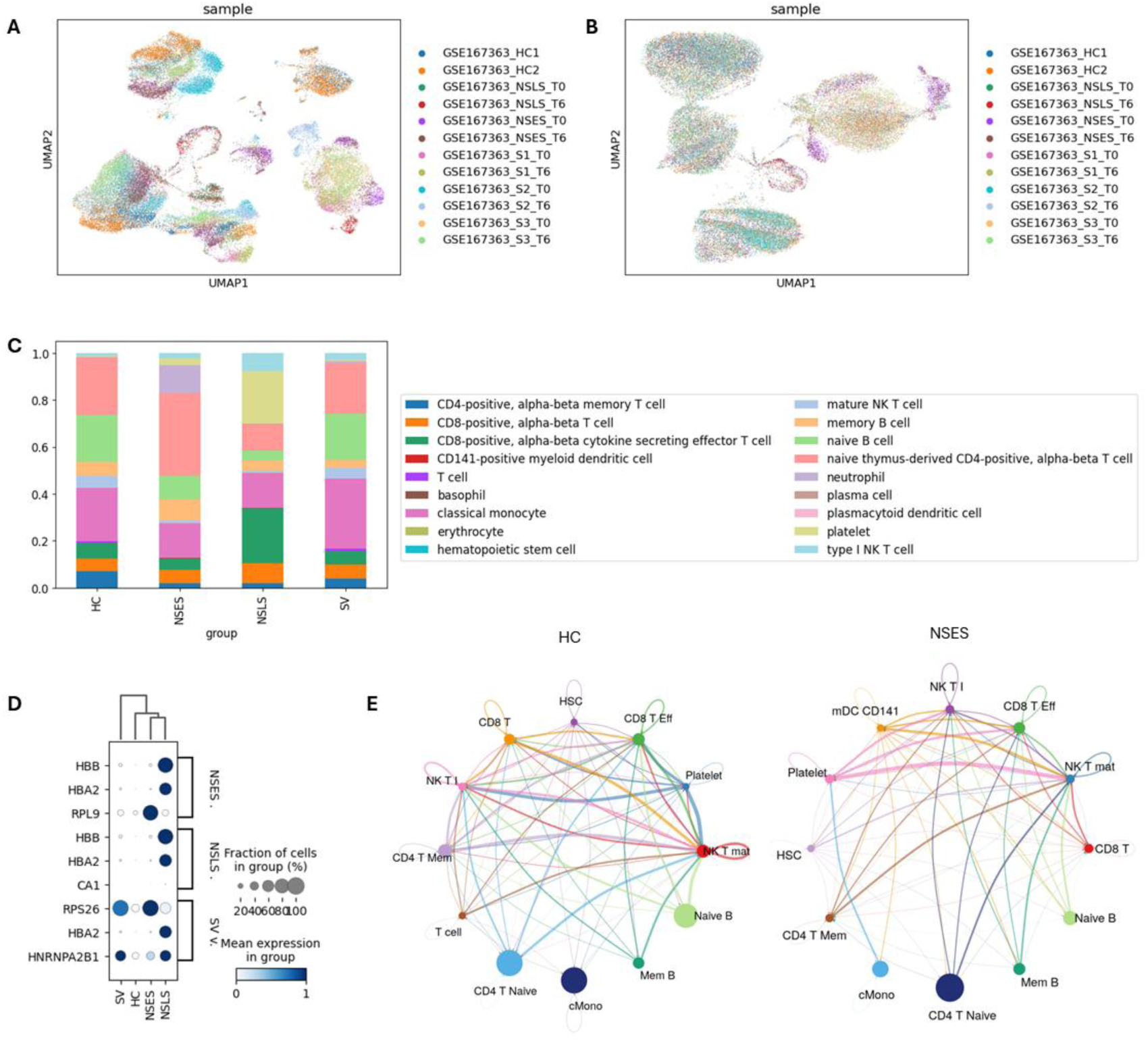
EISCA analysis results on GSE167363 dataset. **A.** UMAP projection of filtered, normalized counts, colored by sample. **B.** UMAP projection of Harmony-integrated counts, colored by sample. **C.** Stacked bar plot comparing predicted cell-type proportions across four groups using scvi-tools and a pre-trained PBMC scANVI model. The stacked bar plot reveals marked immune remodeling in sepsis relative to HC, with a relative expansion of innate compartments (particularly monocytes) and contraction of adaptive lymphocytes, most pronounced in non-survivors. **D.** Dot plot showing differential expression between three sepsis groups and the healthy group. It shows strong expression of hemoglobin genes (HBB, HBA2 and CA1) in the sepsis groups, particularly in the non-survivor groups, which are associated with erythrocyte function and oxygen transport. **E.** Circle plots from the cell–cell communication analysis summarize the overall cell–cell communication network by aggregating communication probabilities between cell types, providing an overview of intercellular interactions.

In the tertiary phase, scvi-tools was applied for automated cell-type annotation using the model *tabula-sapiens-blood-scanvi* from the Hugging Face Model Hub (Figure 6C). The results identified canonical PBMC populations, including CD4⁺ and CD8⁺ T cells, NK cells, B cells, classical and non-classical monocytes, and dendritic cells. The cell-type proportions predicted by EISCA showed strong concordance with those reported in the original study across most clinical groups, with an overall Pearson correlation coefficient of 0.86 (Figure S3). Several of the principal biological observations reported in the original study were reproduced by EISCA. For example, EISCA captured the progressive depletion of B cells with increasing sepsis severity. B cells comprised approximately 26% of cells in HC, 23% in SV, 19% in NSES, and only 9% in NSLS. These estimates closely matched those reported in the original study (approximately 25%, 21%, 16%, and 6% for HC, SV, NSES, and NSLS, respectively). This progressive loss of circulating B cells is consistent with sepsis-associated lymphopenia and was most pronounced in fatal late-stage disease. In addition, EISCA identified marked platelet enrichment in NSLS, supporting the original conclusion that platelet accumulation and dysfunction are associated with fatal sepsis. Some discrepancies were observed for the NSLS group. These differences likely reflect the use of reference-based scANVI label transfer in EISCA compared with the dataset-specific consensus annotation adopted in the original study, as rare or disease-specific cell populations may be underrepresented in the reference dataset.

Differential expression analyses (both cluster-wise and group-wise) were performed using scvi-tools, which identified statistically significant transcriptional programs consistent with systemic inflammation and immune dysregulation. Monocyte clusters in NSLS and NSES showed upregulation of inflammatory and interferon-stimulated genes alongside stress and metabolic pathways. The dot plot shown in Figure 6D highlights differential expression patterns between sepsis groups and healthy controls at cell-level. Hemoglobin genes (HBB and HBA2) and erythroid-associated genes such as CA1 show increased expression in sepsis samples, particularly in non-survivor groups, indicating the emergence of circulating erythroid precursor cells. This observation is consistent with the original study, which reported expansion of erythroid precursors driven by hypoxic stress during severe sepsis. Additionally, changes in ribosomal and RNA-processing genes (RPL9, RPS26, and HNRNPA2B1) suggest altered translational activity and cellular stress responses in immune cells during disease progression.

Cell-cell communication inference further supported these findings, indicating strengthened pro-inflammatory signaling networks involving myeloid populations in non-survivors and comparatively balanced signaling in survivors (Figure 6E). Overall, the EISCA provides a rigorous and statistically grounded characterization of sepsis-associated immune heterogeneity, reproducing key findings from the original publication, including impaired adaptive immunity, platelet-associated innate dysregulation, and distinct immune-cell compositions between survivors and non-survivors.

## 5 Conclusion

In this work, we present EISCA and EISTA pipelines for single-cell and spatial transcriptomics analysis that address the increasing complexity and scale of modern omics data. EISCA provides a comprehensive workflow for scRNA-seq, supporting both droplet- and plate-based technologies, while EISTA extends these capabilities to imaging-based spatial transcriptomics, enabling the integration of molecular and spatial information within tissue contexts. Across primary, secondary, and tertiary analysis phases, both pipelines deliver robust functionalities including quality control, preprocessing, clustering, integration, annotation, differential expression, and cell–cell interaction analysis, with EISTA further incorporating spatial statistical analyses. A key strength of these pipelines lies in their balance between standardization and flexibility. Built on the Nextflow nf-core framework, the pipelines were implemented as modular, scalable, and reproducible workflows that can be executed across local, HPC, and cloud environments without manual dependency management. At the same time, users retain full control over analytical strategies that the workflows can be run either end-to-end or at the level of individual modules, allowing exploratory data analysis with parameter tuning and iterative re-analysis.

From a usability perspective, EISCA and EISTA lower the barrier to entry for non-expert users. Both pipelines provide rapid initial results alongside an out-of-the-box report, enabling immediate assessment of the data and providing a structured foundation for downstream, hypothesis-driven analyses. Their design reflects the inherently dynamic and exploratory nature of single-cell and spatial transcriptomics research. Overall, EISCA and EISTA provide standardized, efficient, and flexible solutions for single-cell and spatial transcriptomics analysis, facilitating reproducible workflows while enabling users to explore complex biological systems at scale.

## 6 Acknowledgements

This work was supported by the Biotechnology and Biological Sciences Research Council (BBSRC), part of UK Research and Innovation; Earlham Institute Strategic Programme Grant Cellular Genomics BBX011070/1 of work package 1: BBS/E/ER/230001A (CellGen WP1 Data Science for Cellular Genomics). We would like to acknowledge Tatsuya Nobori (The Sainsbury Laboratory, Norwich Research Park, Norwich, UK) for providing the spatial transcriptomics datasets and for his valuable support throughout the data analysis and manuscript preparation. We would also like to acknowledge the Galaxy Europe team for their contribution in implementing a Galaxy-based version of the EISTA pipeline, enabling users to apply the pipeline within the Galaxy environment.

## Availability and Implementation

EISCA pipeline is freely available at https://github.com/EarlhamInst/eisca, and EISTA pipeline is freely available at https://github.com/EarlhamInst/eista. A Galaxy workflow implementation of the EISTA pipeline is also freely available at WorkflowHub https://doi.org/10.48546/workflowhub.workflow.2174.5. A hands-on tutorial on using the EISCA pipeline for single-cell RNA-seq analysis is available at https://github.com/Papatheodorou-Group/EISCA_training_course.

## 8 Supplementary materials

**Figure S1.**
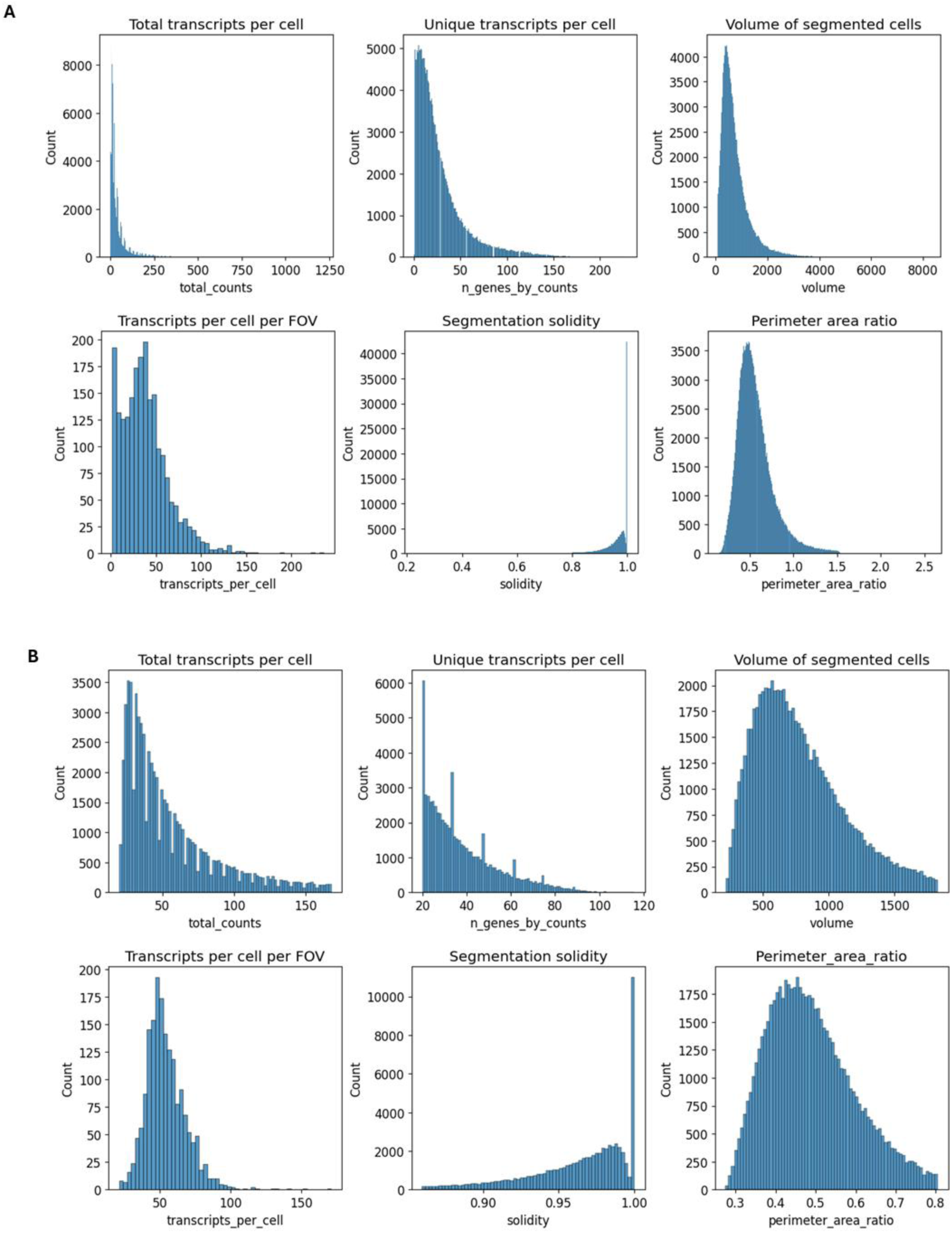
This figure compares histograms of six metrics before and after cell filtering for a Vizgen MERFISH dataset of *Arabidopsis thaliana* leaves from the 24 h sample. **A.** Histograms of the distributions of six metrics from the raw counts data. **B.** Histograms of the distributions of the same six metrics after cell filtering based on count distributions and segmentation quality metrics implemented in EISTA, including minimum total counts, upper bound for total counts, minimum number of detected genes, upper bound for cell volume, lower bound for transcripts per cell per FOV, lower bound for solidity, and upper bound for the perimeter-to-area ratio.

**Figure S2.**
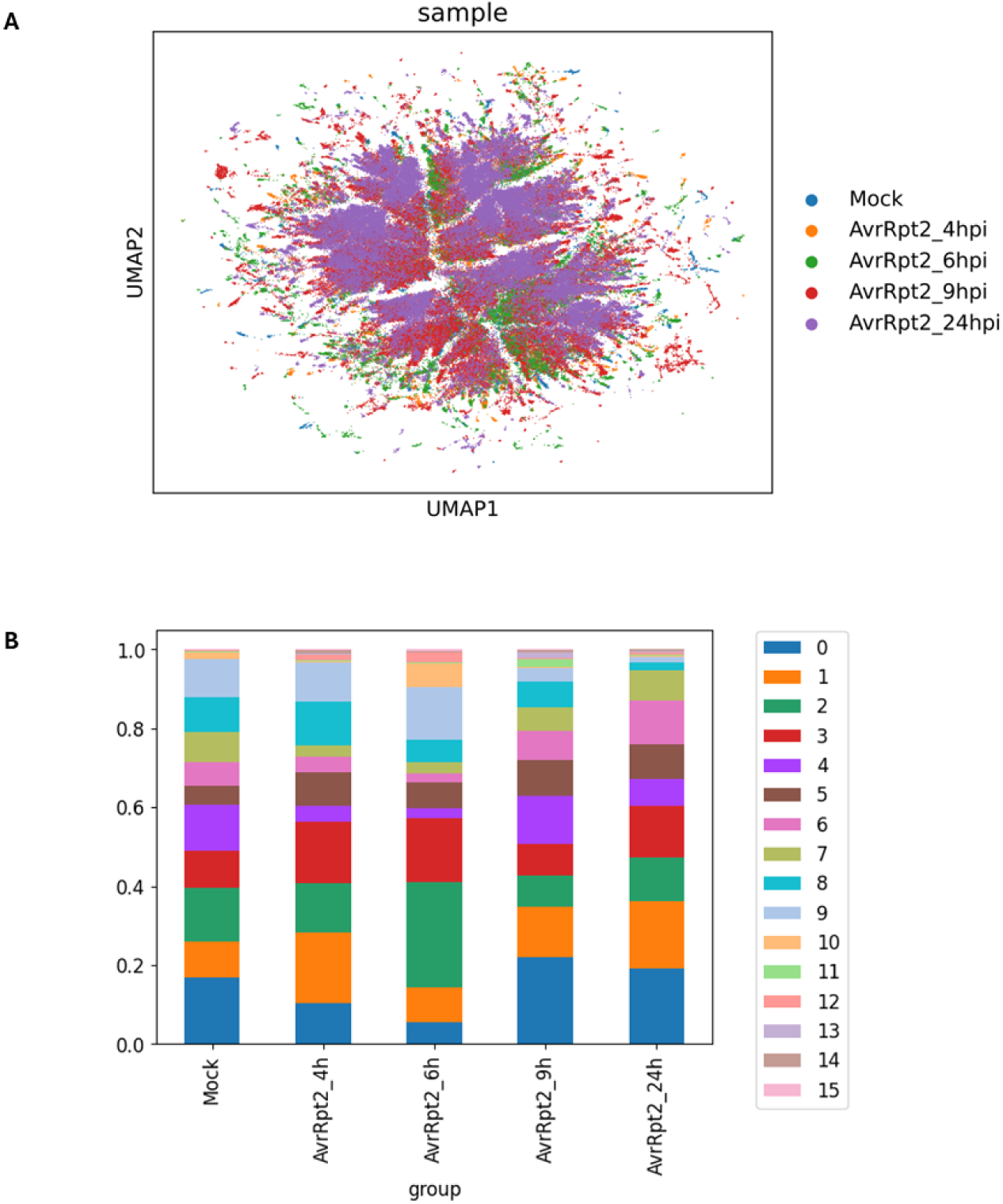
This figure shows clustering results using Harmony integration for a Vizgen MERFISH dataset of *Arabidopsis thaliana* leaves. **A.** UMAP projection of clustering with Harmony integration, colored by sample. Compared to the clustering results without integration (Figure 4B), the Harmony-integrated UMAP shows a starfish-like pattern, suggesting potential over-correction and partial removal of the biological trajectory related to infection dynamics. Therefore, we use the clustering results without integration for the case study, as they better preserve the underlying biological signal. **B.** The corresponding stacked bar plot shows cluster proportions at resolution 0.2 after Harmony integration. We can see that all samples have similar cluster composition profiles, rather than distinct condition-specific profiles (Figure 4D), suggesting potential overcorrection.

**Figure S3.**
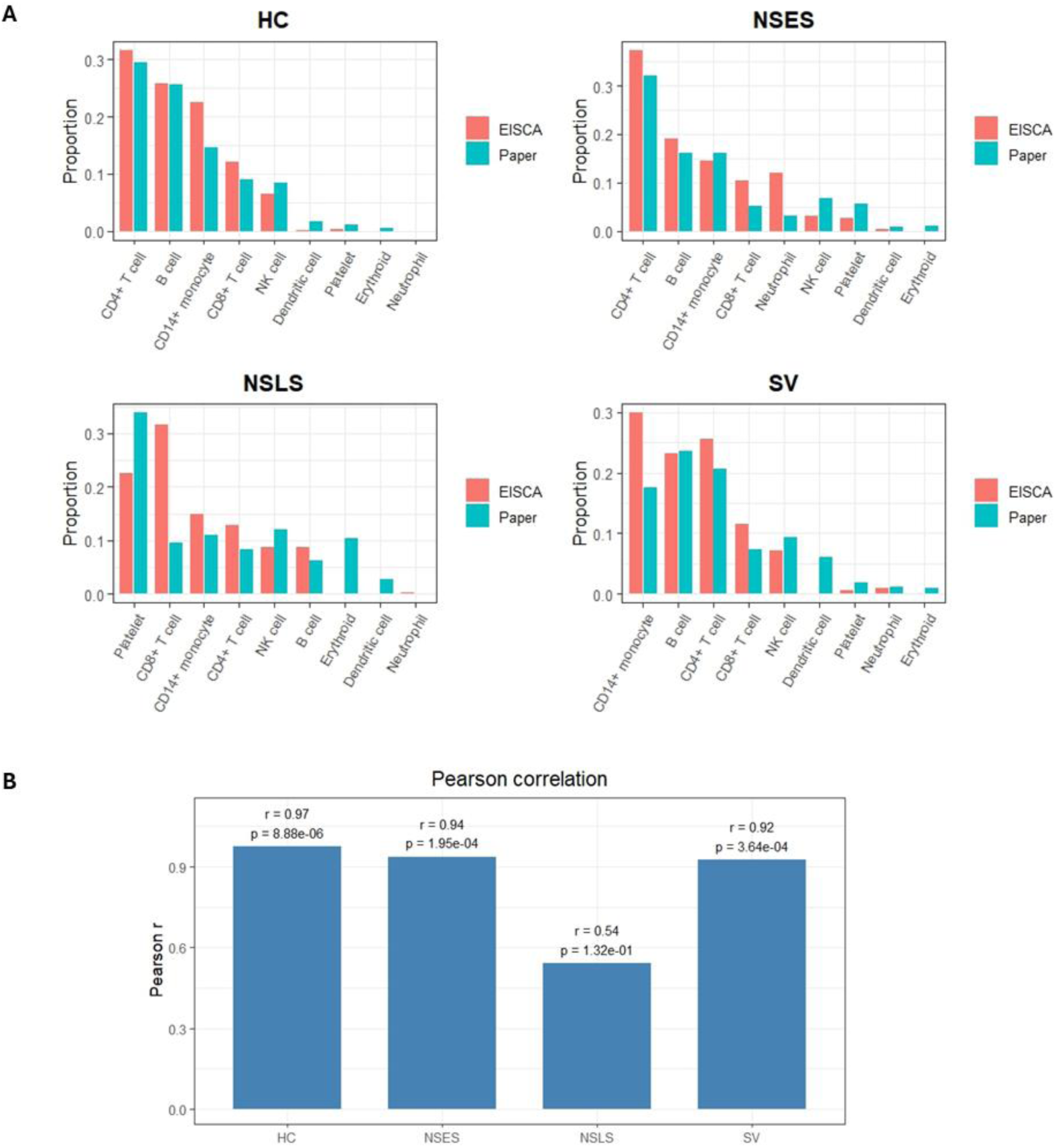
Comparison of predicted cell types between the scVI-based analysis in the EISCA pipeline and the original published results (“Paper”) for the scRNA-seq dataset from a human sepsis study (Qiu et al., 2021). **A.** Bar plots comparing the estimated cell-type proportions obtained by EISCA and the original paper for healthy controls (HC), sepsis survivors (SV), and non-survivors at the early (NSES) and late (NSLS) stages of disease. To enable comparison between the two sets of results, the predicted cell types from the pre-trained PBMC scANVI model were harmonized as follows: B cell (naive B + memory B + plasma cell), CD4+ T cell (CD4-memory + naive thymus-derived CD4), CD8+ T cell (conventional CD8 + cytokine-secreting effector CD8), CD14+ monocyte (classical monocyte), Dendritic cell (CD141-positive myeloid DC + plasmacytoid DC), NK cell (mature NK T + type I NK T), Erythroid (erythrocyte). **B.** Bar plot showing the Pearson correlation coefficients between the cell-type proportions estimated by EISCA and those reported in the original paper across the four study groups. P-values from the Pearson correlation tests are also shown.

